# Person-specific intrinsic-timescale topographies persist from rest into sentence processing

**DOI:** 10.64898/2026.09.28.755139

**Authors:** Daniel Di Giovanni, Ida Toivonen, Georg Northoff

## Abstract

Language processing does not occur in a vacuum. Language comprehension and production have its own temporal organization, and it unfolds against ongoing neural. We examined whether the base organization persists from rest into sentence processing, and we further examined how it relates to evoked responses and semantic decisions. Sentences crossed linguistic argument versus adjunct completions with semantic plausibility. Anomalies elicited a centroparietal late positivity from 600 to 900 ms (+0.628 µV, 95% CI 0.349–0.907). Channel-wise autocorrelation timescales formed person-specific topographies: own-person rest–task similarity exceeded mismatched-person similarity (Fisher-z advantage = 0.463, permutation P = 5.00 × 10^−6^), identifying 78.6% of participants against 7.1% chance. The advantage survived Cz re-referencing and surface-Laplacian transformation. In separate ERP predictor tests, longer absolute task timescales accompanied a smaller anomaly effect (cross-fitted Spearman ρ = −0.486, Holm-adjusted P = 0.030). Stronger topographic continuity also accompanied greater anomaly-related decision slowing, although the behavioral association fell short of correction across six endpoints (ρ = 0.547, familywise P = 0.054). Supporting analyses evaluated larger eligible samples and an eyes-closed subset. The findings place persistent, person-specific temporal organization within sentence processing and link absolute task timescale to the neural response to semantic anomaly.

## Introduction

Stimulus-evoked activity emerges within an ongoing neural state. Its interpretation therefore depends not only on the response to an experimental event, but also on the temporal organization of the activity in which that response occurs [1]. Spatiotemporal accounts of brain function propose that intrinsic dynamics provide a context for integrating incoming information [2–5]. A central question is whether this organization remains identifiable as people move from rest into a cognitive task, and whether it relates to their responses within that task.

Intrinsic neural timescales quantify how long activity remains correlated with itself. These timescales differ across cortical systems, forming a hierarchy of temporal integration capacities [6,7]. Longer timescales can support integration over extended contexts, whereas shorter timescales allow more rapid differentiation [2,3]. Their organization combines stability across states with sensitivity to behavioral demands [2,3,8]. This makes intrinsic timescales a useful candidate for linking ongoing activity to the processing of temporally structured input.

Distributed brain patterns can preserve individual identity across cognitive states, and task-free functional organization can predict aspects of task-evoked activity [9–11]. These findings motivate two complementary measurements. The spatial arrangement of timescales can reveal whether a participant retains a characteristic configuration from rest into task. The absolute duration of those timescales can then be related to an evoked response. Neither measurement alone establishes how the configuration becomes relevant to a defined cognitive operation.

Spoken language processing offers such an operation because the processing of a sentance requires information to be accumulated across words and related to an event. Verb arguments are selected by a predicate and typically denote core event participants; adjuncts are less tightly selected and often contribute manner, time, or place information [12,13]. For example, consider this sentence: *Last week, the manager sent a file to Sara over email*. In this sentence, *the manager, a file*, and *to Sara* are arguments, whereas *last week* and *over email* are adjuncts. Argumenthood is graded across syntactic, semantic, and discourse dimensions [12–17]. The argument–adjunct contrast therefore allows us to examine semantic decisions made under different degrees of lexical constraint, without assuming that the two completion types must have different mean neural timescales.

We analyzed eyes-open resting EEG and EEG during a spoken-sentence task crossing argument versus adjunct completions with semantic plausibility. Participants first indicated that they had understood each sentence, then judged whether its subject had been affected. Verified critical-word offsets aligned the EEG to the completion word.

The primary neural endpoint was preservation of the person-specific intrinsic-timescale topography from rest into task. We then examined two functional relationships, first, absolute resting and task timescales as separate predictors of the late-positive ERP anomaly effect, and topographic continuity with anomaly-related semantic decision slowing. Cross-fitting separated task observations used for predictor and outcome in both directions. Additional cohorts and control analyses provide supporting evidence in the Supplementary Materials.

## Methods

### Participants and ethics

The study cohort comprised of an archive of 22 healthy, right-handed, native monolingual English-speaking adults. The main analysis cohort comprised the 14 participants with eligible eyes-open rest EEG, sentence-task timescale estimates, all-block and split-half ERP effects, and usable behavior. Eligibility followed the fixed recording, artifact, and trial-coverage criteria described below; missing participant-level measurements were not imputed. The experimental protocol was approved by the research ethics committee of the University of Ottawa Institute of Mental Health Research at The Royal, Ottawa, Canada. All participants provided written informed consent before participation.

### Sentence task

Participants heard 640 trials in eight blocks of 80. The 2 × 2 target design crossed completion structure (argument, adjunct) and semantic plausibility (plausible, anomalous), with 128 trials per condition. Another 128 bare-verb trials were excluded from these language contrasts. Four verbs (*ate, cooked, drank*, and *packed*) occurred with the four completion types. For example, *The student ate yogurt* and *The student ate baggage* contained plausible and anomalous arguments; *The student ate quickly* and *The student ate tightly* contained plausible and anomalous adjuncts. The 16 target utterances and four bare-verb utterances were repeated across blocks.

After each sentence, participants pressed a button to indicate that they had understood it, then judged whether the subject had been affected. The affected-subject response probed event-participant interpretation; the understood response provided a comparison stage. Reaction times were retained for both responses.

Critical-word onset was the task stimulus marker plus the sentence-specific acoustic offset verified from the original audio files. Marker codes, E-Prime timing, offsets, and recording bounds were checked trial by trial. Trials without a valid target-condition label or an available EEG epoch were excluded.

### EEG acquisition

The acquisition protocol used a 64-channel Ag/AgCl cap, a Neuroscan amplifier, DC recording at 1,000 Hz, a nose-tip online reference, EOG channels, electrode impedances below 5 kΩ, and a 60-Hz notch. A standard 10–20 montage was assigned; M1, M2, CB1, and CB2 were treated as auxiliary channels, leaving 60 scalp channels for analysis.

### Preprocessing and quality control

Continuous data were processed with zero-double-phase, fourth-order Butterworth filters in three branches so that ERP filtering did not determine timescale estimates: 0.1–45 Hz for dynamics, 0.5–40 Hz for conventional ERP data, and 1–40 Hz for diagnostic channel assessment. The final dynamics and ERP branches were re-referenced to the mean of the good scalp channels and resampled to 250 Hz. On the diagnostic copy, a channel was flagged if its SD was <0.2 µV, if its robust log-SD z score exceeded 6 together with median absolute correlation <0.30 (or exceeded 10 regardless of correlation), or if its median absolute correlation with the 2-s-window scalp median was <0.30. A recording failed if >20% of scalp channels were flagged. FastICA used at most 30 components (decimation factor 4); up to eight components associated with HEO/VEO or ECG were removed. Flagged channels in retained recordings were interpolated by spherical splines before resampling.

For fixed artifact acceptance, task data were segmented from −1 to +3 s relative to the verified critical-word onset and rest was divided into non-overlapping 4-s segments. A segment was rejected if any retained scalp channel contained nonfinite values, exceeded 200 µV peak-to-peak, or had SD <0.1 µV. Rest required at least 120 s of accepted data; each block × verb × structure × plausibility cell required at least four accepted task trials. These thresholds and recording-level bad-channel decisions were fixed before inferential analysis.

### Intrinsic timescales

We quantified intrinsic timescales using autocorrelation-window measures [2,3,18]. For each accepted 4-s segment or trial and channel, we demeaned the signal and computed an unbiased autocorrelation by dividing each autocovariance lag by its number of overlapping sample pairs, then normalizing by zero-lag covariance. The autocorrelation-window zero crossing (ACW-0) was the first lag with autocorrelation ≤0 within a maximum evaluated lag of 2 s; estimates without a crossing were recorded as missing rather than capped. At 250 Hz, temporal resolution was 4 ms. Estimates were obtained within segment boundaries and summarized by the median, avoiding artificial correlations across concatenated segments.

For each participant, the resting topography was the median timescale across accepted rest segments at each of 60 scalp channels. Task estimates were first summarized within block × verb × structure × plausibility conditions, then averaged within channel with weights proportional to accepted trial counts. Only the four target conditions entered. Pearson correlations across corresponding channels formed a 14 × 14 task–rest similarity matrix and were converted to Fisher z; at least 20 jointly finite channels were required for each pairing. The continuity score was each participant’s own-person similarity minus their mean similarity to the other 13 resting topographies. Identity was assigned to the rest participant with maximal similarity. All-block and split-half continuity scores used this same 14-person reference pool. We tested the mean own-person advantage and identity accuracy with 200,000 participant-label permutations.

### Reference and surface-Laplacian sensitivity

Reference sensitivity was assessed by recomputing timescales from Cz-referenced and surface-Laplacian-transformed EEG. Cz was omitted from Cz-reference spatial correlations because its signal becomes identically zero. Current-source density (CSD) used spherical splines with an automatically fitted sphere, λ^2^ = 10^−5^, stiffness = 4, and 50 Legendre terms [19,20]. The existing average-reference trial and segment acceptance decisions were inherited unchanged. The behavioral continuity score and absolute-timescale ERP predictors used CSD timescales; the ERP outcome used average-reference voltage.

### Behavioral coefficients

Reaction times from 100 to 10,000 ms were log transformed. For each participant and task split, ordinary least squares estimated coefficients for anomaly (anomalous = +0.5, plausible = −0.5), structure (argument = +0.5, adjunct = −0.5), their interaction, standardized chronological block order, and verb indicators. Thus, the structure coefficient was the adjusted argument-minus-adjunct log-time contrast. Percentage differences were 100 × [exp(β) − 1]. Group contrasts used one-sample t-tests. Alternative response-time estimators and condition decompositions are reported in the supplement.

### Cross-fitting and multiplicity in the behavioral analysis

Within each participant, the four blocks of each argument or adjunct family were divided into positions 1 and 3 versus 2 and 4. Both halves contained both completion types. We correlated odd-half continuity with the even-half behavioral coefficient, then reversed the pairing and averaged the two Spearman correlations. Both correlations used the same 14 participants, with each task-half topography compared against their full resting topographies. A joint participant-label permutation moved both behavioral halves together. Cross-fitting separated task observations, rather than dividing participants into smaller samples.

The behavioral multiplicity family retained six endpoints: anomaly, structure, and interaction coefficients for the affected-subject and understood responses. Westfall–Young single-step minP correction used 200,000 joint participant-label permutations; Holm-adjusted values are also reported in Supplementary Table S3. The anomaly coefficient was the focal behavioral endpoint, and the structure-by-anomaly coefficient tested moderation. This family was retained when the study cohort was restricted to 14 participants. Leave-one-participant-out estimates and alternative estimators assessed sensitivity.

### Supporting cohorts and uncertainty

The supplementary analyses retain the larger samples eligible for behavior (N = 21), the group ERP effect (N = 18), and eyes-open rest–task matching (N = 15). The 11 main-cohort participants with both resting states provide a matched eyes-closed comparison; the full eyes-closed sample contained 13. These samples support the main cohort analysis and are labeled separately throughout the supplement. Behavioral estimates based on the 15-person resting reference pool are also presented there. Pointwise 95% association intervals used 5,000 paired participant percentile-bootstrap resamples with both halves carried together and the reference pool held fixed. Pearson and Spearman split-half correlations assessed measurement reproducibility.

### ERP analysis

ERP data were low-pass filtered at 30 Hz and epoched from −200 to 1,000 ms relative to critical-word onset, with baseline correction from −200 to 0 ms. Trial acceptance was inherited from the corresponding 4-s artifact window. The endpoint was mean 600–900-ms amplitude over C1, Cz, C2, CP1, CPz, CP2, P1, Pz, and P2. Participant effects were anomalous minus plausible, pooling argument and adjunct trials. Eligibility required at least 80 accepted trials per plausibility level overall and 40 per level in each task half. Every presented participant met both requirements. Response-timing and structure-specific sensitivities are reported in the supplement.

Two absolute timescales were tested separately as ERP predictors. Resting timescale was the median across channels of their full clean eyes-open resting estimates, in milliseconds. Task timescale used accepted-trial-weighted condition means within channel, then the median across channels. The same finite channel intersection across rest and all/odd/even task estimates was used within each participant, with at least 20 channels required. Absolute timescales were not spatially centered and did not use mismatch subtraction.

The resting test correlated the full-rest timescale with the all-block ERP anomaly effect. The task test averaged the correlations pairing odd-half task timescale with even-half ERP and vice versa. Both directions retained all participants. Two-sided inference used 200,000 participant-label permutations, moving ERP columns jointly, and Holm correction across the resting and task tests. Pointwise intervals used 5,000 paired participant bootstrap resamples. The predictors were evaluated separately, without mutual adjustment or validation in new participants.

### Statistical reporting

Participants were the sampling units for brain–behavior and ERP associations [21–23]. Tests were two-sided except for directional own-person matching and identity tests. Permutation P values used (exceedances + 1)/(permutations + 1). Group-mean intervals were 95% t intervals, and Cohen’s *d*_*z*_ was the participant-level mean divided by its SD. Behavioral and absolute-timescale ERP tests formed separate secondary multiplicity families; their corrections do not cover every analysis in the study. Further methods for condition selectivity and temporal variability accompany their results in the supplement.

## Results

### Anomalies elicited a group late positivity

In the study cohort, anomalous sentences elicited a centroparietal late positivity from 600 to 900 ms (Fig. 1C,D). The pooled anomalous-minus-plausible effect was +0.628 µV (95% CI 0.349–0.907; dz = 1.30; P = 3.07 × 10^−4^). The larger ERP-eligible sample supported this group response (Supplementary Section S5).

**Figure 1.**
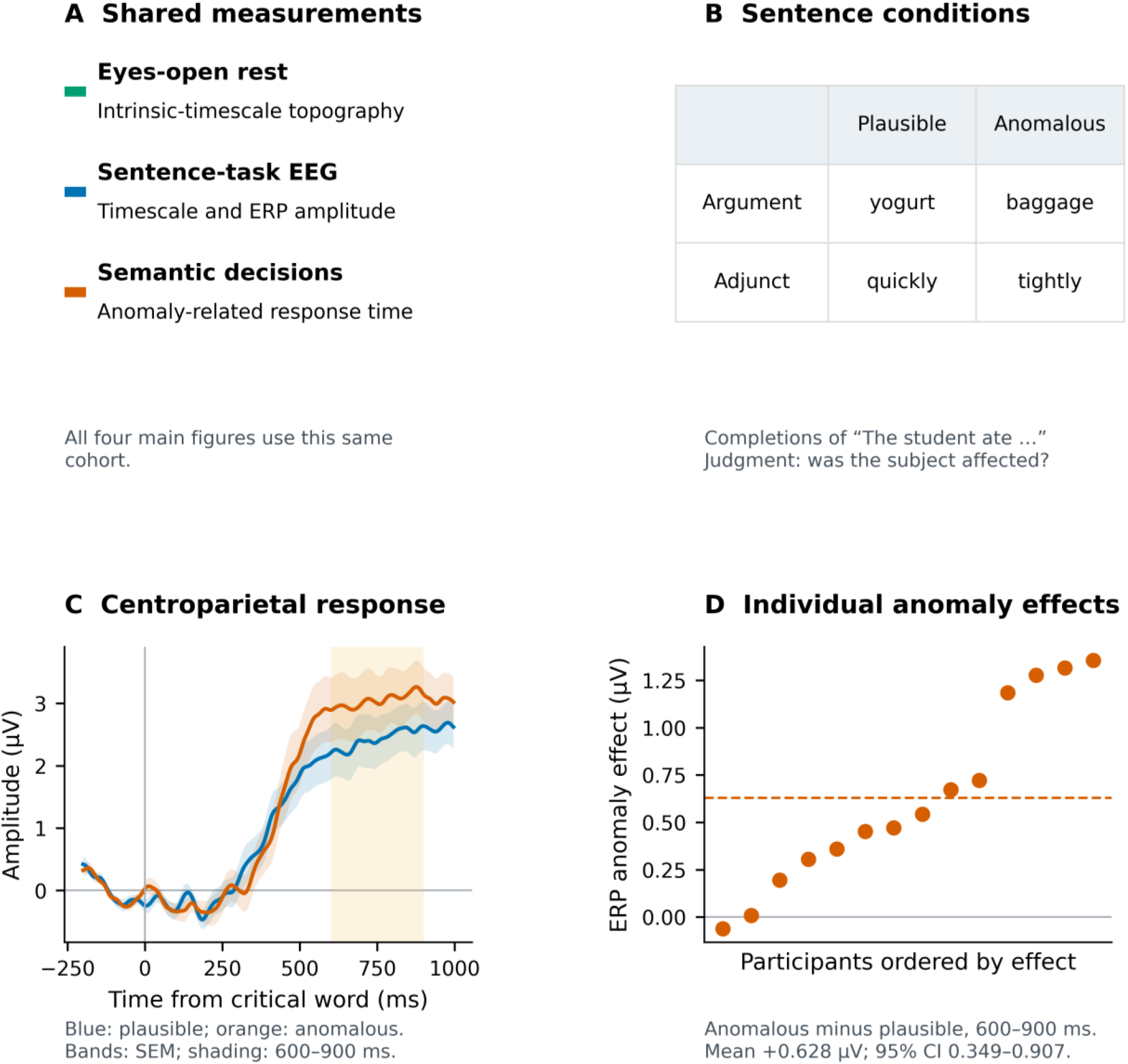
Sentence processing. All panels refer to the participants with eyes-open resting EEG, task EEG, ERP effects, and behavior. (A) Measurements available in every participant. (B) Example argument and adjunct completions crossed with plausibility. (C) Average-reference centroparietal waveforms, with participant mean ± SEM; shading marks 600–900 ms. (D) Individual anomalous-minus-plausible effects, ordered by amplitude; the dashed line marks the group mean.

The group response provides a physiological anchor for the timescale analysis, but its individual amplitude contrast had limited split-half reliability (Pearson r = 0.206; Spearman ρ = 0.156). In these same participants, 59.6% of retained trial response markers preceded 900 ms. We therefore describe a late positivity in the P600 window, allowing for decision and motor contributions (Supplementary Fig. S2).

### Timescale topographies preserve participant identity from rest into language

Rest–task timescale topographies preserved participant identity in the study cohort (Fig. 2). Average-reference own-person similarity was 0.628 in Fisher-z units, compared with a mean mismatched similarity of 0.164. The own-person advantage was 0.463 (95% CI 0.340–0.587; one-sided permutation P = 5.00 × 10^−6^). Eleven of the 14 task topographies were matched to their own resting recording: 78.6% identification against 7.1% chance (permutation P = 5.00 × 10^−6^).

**Figure 2.**
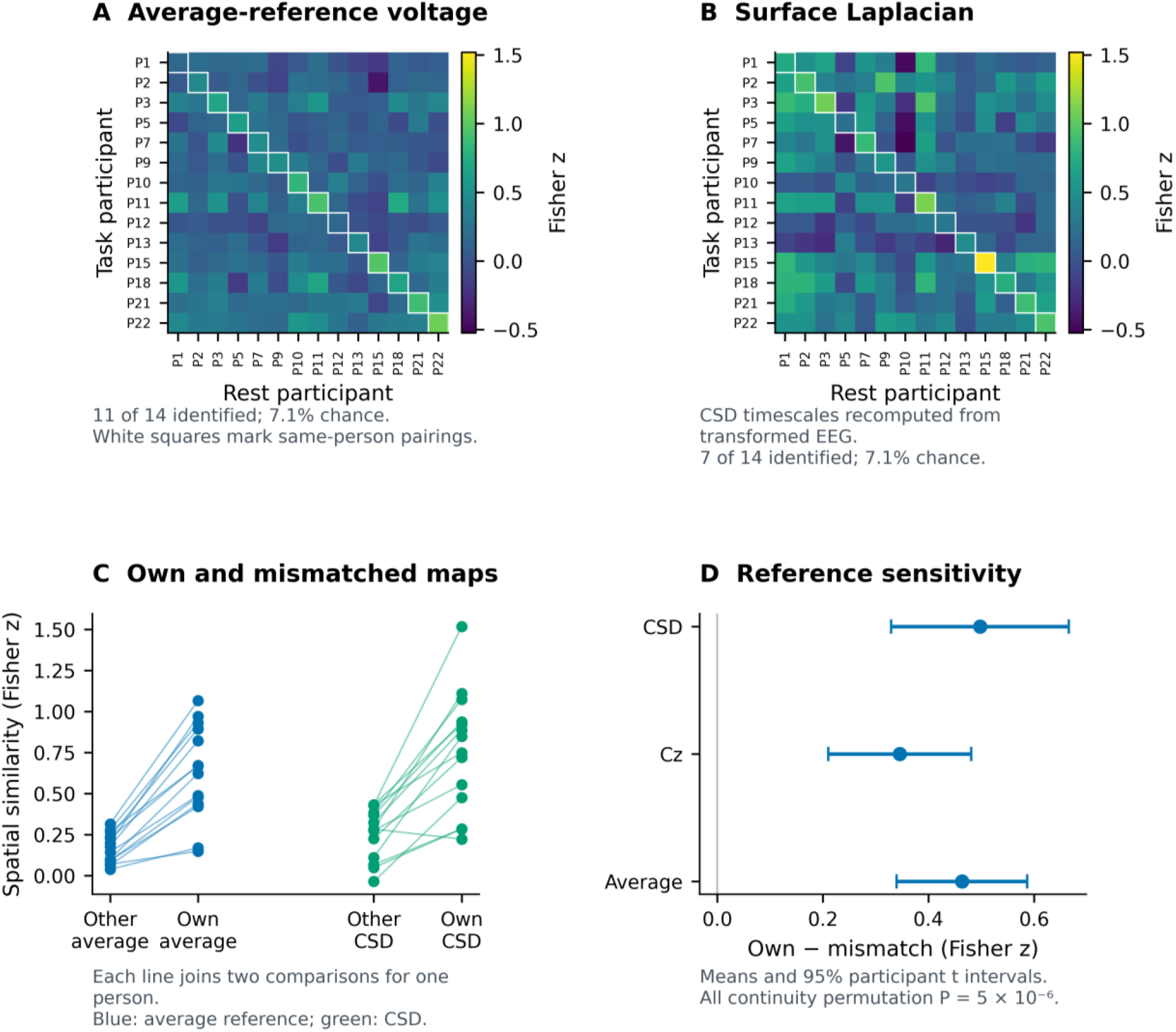
Person-specific timescale topographies. (A) Average-reference and (B) CSD task–rest similarities in Fisher-z units; diagonal squares compare a participant with their own rest recording. (C) Paired own-person and mean mismatched-person similarities. (D) Mean own-person advantages with 95% participant-level t intervals. Above-chance identification and continuity tests use participant-label permutations.

The advantage persisted after Cz re-referencing (Δz = 0.346, 95% CI 0.210–0.481) and surface-Laplacian transformation (Δz = 0.498, 95% CI 0.329–0.666; both continuity permutation P = 5.00 × 10^−6^). Each transformation identified seven of 14 participants, above the same 7.1% chance level. Thus, preservation of the timescale pattern was not confined to one reference scheme.

Supporting analyses preserved the fingerprint in the 11 participants with both resting states and after group-centering and exclusion of pre-interpolation bad channels in the larger 15-person eyes-open sample. These controls constrain explanations based on a common scalp pattern or flagged channels, while retaining the limitations of a same-session, same-cap comparison (Supplementary Section S2 and Fig. S1).

### Longer absolute task timescales were associated with smaller ERP anomaly effects

Absolute task timescale was negatively associated with the late-positive anomaly effect in the main cohort (mean cross-fitted Spearman ρ = −0.486, 95% CI −0.741 to −0.074; participant-label permutation P = 0.0151; Holm-adjusted P = 0.0302; Fig. 3). Both directions were negative (odd task to even ERP: ρ = −0.327; reverse: ρ = −0.644), and leave-one-participant-out estimates ranged from −0.566 to −0.398. Longer task timescales accompanied a smaller anomalous-minus-plausible late positivity.

**Figure 3.**
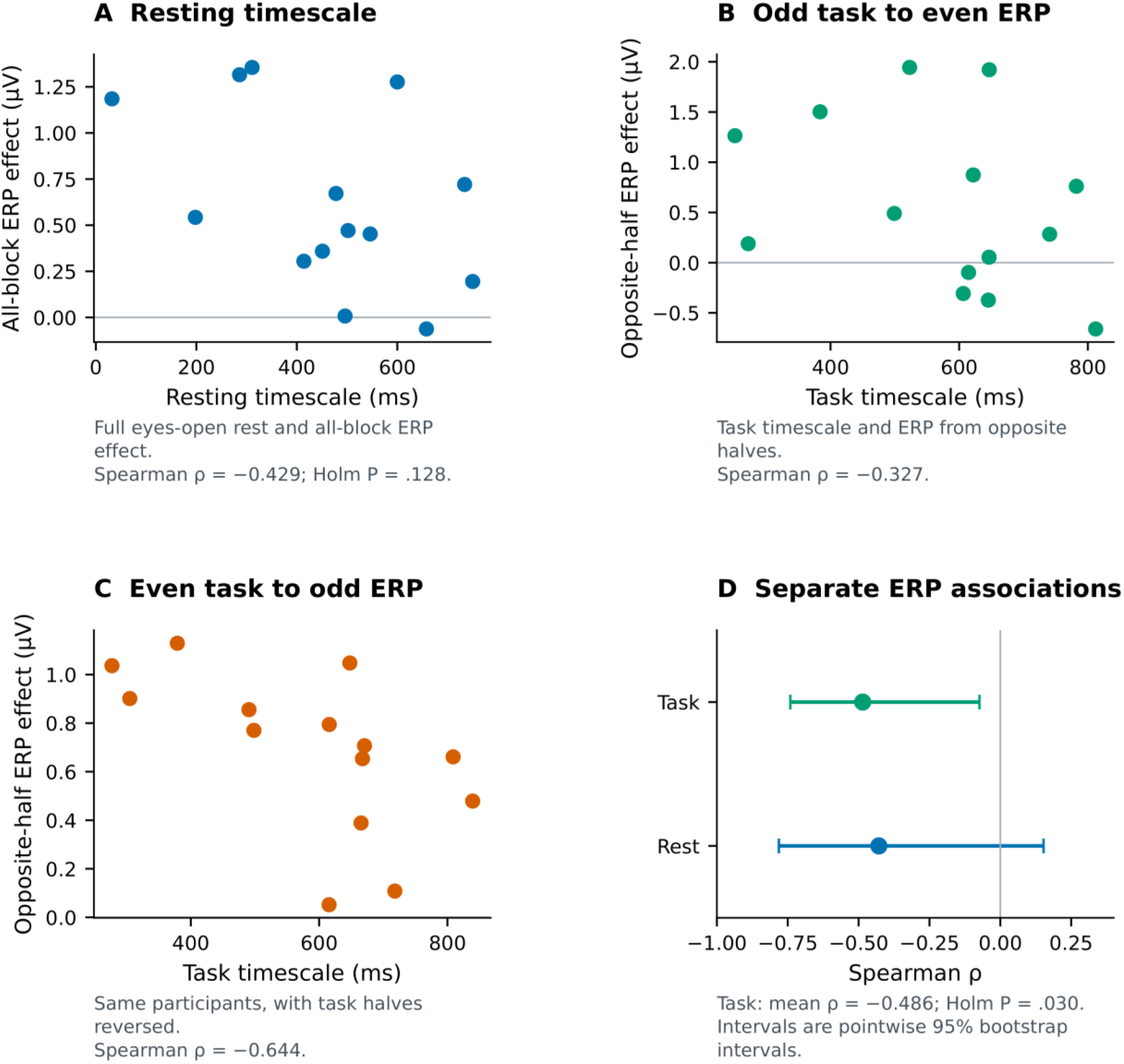
Absolute timescales and the late-positive anomaly response. (A) Full eyes-open resting CSD timescale versus the all-block ERP contrast. (B,C) Task timescale versus the opposite-half ERP contrast, with the task halves reversed between panels. (D) Separate resting and task associations with pointwise 95% participant-bootstrap intervals. The task statistic averages the two cross-fitted correlations. Holm correction covers both predictors (adjusted P = 0.128 for rest and 0.030 for task). Timescales are channel medians in milliseconds; ERP effects are anomalous-minus-plausible average-reference voltage from 600 to 900 ms.

The absolute resting-timescale estimate was negative but inconclusive (ρ = −0.429, 95% CI −0.781 to 0.153; Holm-adjusted P = 0.128). Task timescale itself was reproducible across halves (Pearson r = 0.988; Spearman ρ = 0.925). The resting and task tests are reported together in Table 1; their different significance levels do not establish a difference between states.

**Table 1.** Main analysis results.

| Endpoint | Estimate | 95% interval | Inference |
| --- | --- | --- | --- |
| Average-reference continuity | $\Delta z = 0.463$ | 0.340 to 0.587 | Permutation $P = 5.00 \times 10^{-6}$ |
| CSD continuity | $\Delta z = 0.498$ | 0.329 to 0.666 | Permutation $P = 5.00 \times 10^{-6}$ |
| Participant identification | 11/14 (78.6%) | Chance = 7.1% | Permutation $P = 5.00 \times 10^{-6}$ |
| Late-positive anomaly effect | +0.628 $\mu V$ | 0.349 to 0.907 | $P = 3.07 \times 10^{-4}$ |
| Resting timescale-ERP | $\rho = -0.429$ | -0.781 to 0.153 | Holm $P = 0.128$ |
| Task timescale-ERP | $\rho = -0.486$ | -0.741 to -0.074 | Holm $P = 0.030$ |
| Argument response advantage | 16.8% faster | 8.9% to 24.0% | $P = 7.47 \times 10^{-4}$ |
| Continuity-anomaly cost | $\rho = 0.547$ | 0.197 to 0.764 | Familywise $P = 0.054$ |

$\Delta z$ is own-person minus mean mismatched-person Fisher-z similarity, with the other 13 participants defining the mismatch pool. CSD, current-source density; ERP, event-related potential; $\rho$ , Spearman correlation. The behavioral and task-timescale associations average opposite-half correlations. Intervals are pointwise 95% t intervals for means and percentile-bootstrap intervals for associations; they are not multiplicity-adjusted. The behavioral correction covers six endpoints and the ERP correction covers the two absolute-timescale predictors. Rest and task were tested separately; their significance levels do not test a between-state difference. Identification chance is 1/14.

### Intrinsic continuity and semantic decision dynamics

Affected-subject judgments were faster following argument than adjunct completions. The adjusted log-time contrast was −0.184 (95% CI −0.274 to −0.093; P = 7.47 × 10^−4^), corresponding to 16.8% faster responses (95% CI 8.9–24.0%; Fig. 4A). The anomaly coefficient varied across participants, providing the individual behavioral contrast for the continuity analysis.

**Figure 4.**
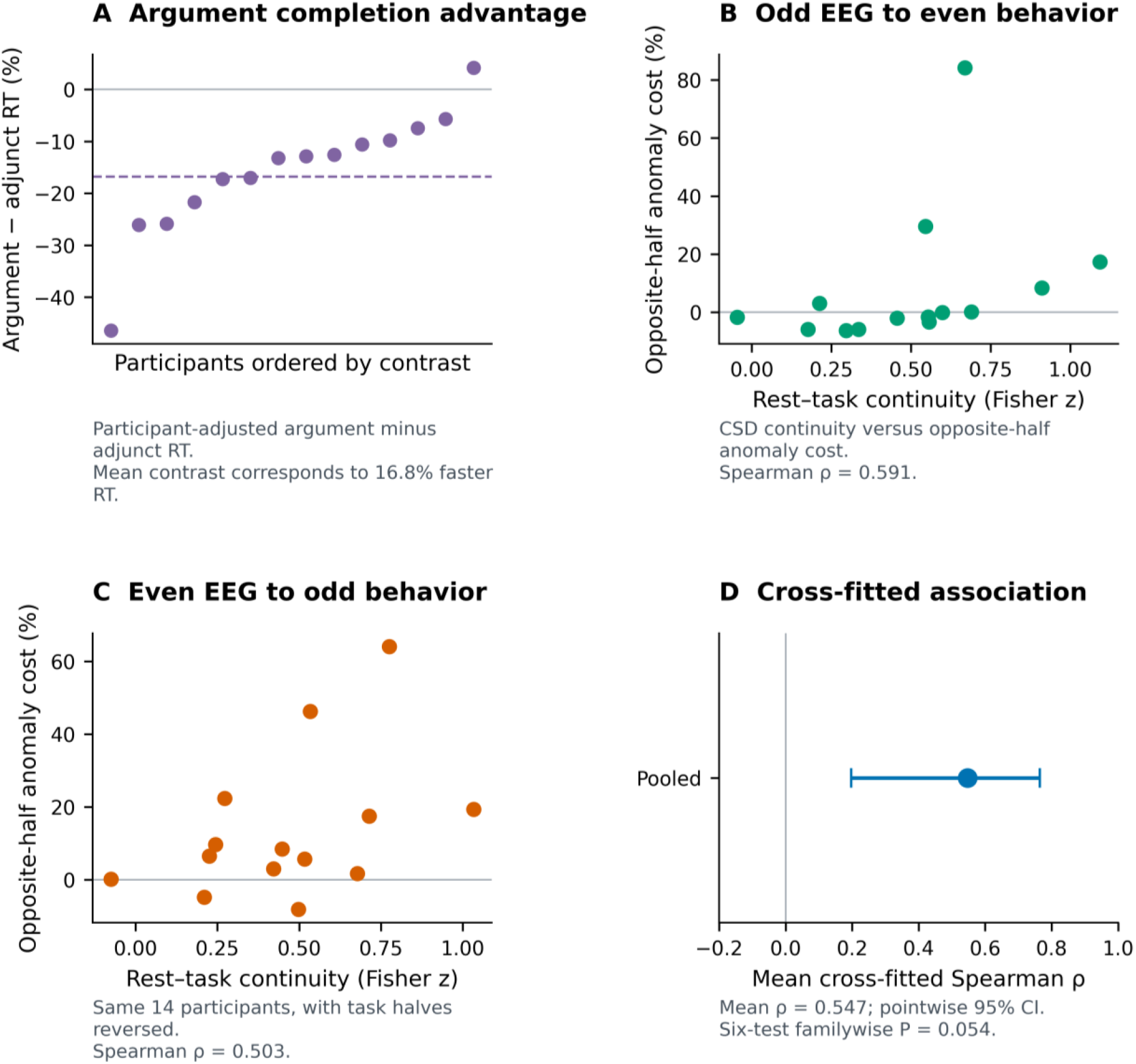
Intrinsic continuity and affected-subject decisions. (A) Adjusted argument-minus-adjunct log-time coefficients converted to percentage differences; the dashed line marks the transformed mean coefficient. Negative values indicate faster argument judgments. (B,C) The two opposite-half continuity–anomaly associations, with anomaly costs converted to percentage change. Both halves retain all participants. (D) Their mean Spearman association with a pointwise 95% participant-bootstrap interval. The six-test familywise P value is 0.054; the positive estimate does not meet that corrected threshold.

Stronger rest–task continuity accompanied greater anomaly-related slowing in both cross-fitting directions (odd continuity to even behavior: ρ = 0.591; reverse: ρ = 0.503; Fig. 4B,C). The mean association was ρ = 0.547 (pointwise 95% CI 0.197–0.764; nominal participant-label permutation P = 0.0093). Correction across all six behavioral endpoints gave familywise P = 0.054 and Holm-adjusted P = 0.056. The positive estimate therefore fell short of the familywise significance threshold (Fig. 4D).

Leave-one-participant-out estimates remained positive (ρ = 0.470–0.596), and alternate response-time estimators gave similar directions. The estimate was sensitive to the resting reference pool, as the analysis using the larger 15-person pool gave ρ = 0.574 and familywise P = 0.031 (Supplementary Section S4). The structure-by-anomaly association did not establish argument–adjunct moderation after correction (familywise P = 0.369; Supplementary Table S3).

## Discussion

A combination of rest EEG, task physiology, and behavior allowed us to examine intrinsic temporal organization at three related levels. Person-specific timescale topographies persisted from rest into sentence processing. Absolute task timescale related to the amplitude of the late-positive anomaly response. Topographic continuity also showed a positive relationship with semantic decision slowing, although that behavioral estimate was sensitive to the reference pool and did not meet the six-test familywise threshold in the analysis.

The fingerprint result fits a spatiotemporal account in which ongoing dynamics provide a temporal context for cognition [1,4,5,24]. A timescale topography describes how temporal persistence is distributed across the scalp. Its preservation across rest and sentence processing suggests that task engagement occurs within a participant-specific configuration, rather than requiring a wholly new temporal organization. The result survives reference changes and is supported by separate-state and channel controls. Work distinguishing spontaneous and evoked dynamics further motivates asking how this persistent organization relates to the activity elicited by a stimulus [25].

The spatial pattern and the absolute timescale address complementary aspects of that organization. Continuity measures preservation of relative channel differences; the absolute predictor summarizes the duration of temporal persistence. This makes it possible to relate a stable configuration to task physiology and behavior without confusing differences in sample composition. The present associations do not establish independent mechanisms or a causal path from resting activity through ERP amplitude to a decision.

ERP research distinguishes semantic retrieval from later integration, reanalysis, and updating [26–29]. Here, the positive group contrast shows a clear mean response in the P600 window, while longer absolute task timescales accompany a smaller anomaly-minus-plausible contrast. Recent work relating intrinsic timescales to event-related activity [30] motivates studying this connection across different tasks and measures. The present analysis concerns a signed EEG language contrast; it does not assign the effect to a specific mechanism. Related work linking neural and behavioral temporal dynamics during music engagement [31] likewise encourages connecting intrinsic organization to well-defined responses, while keeping the temporal measures and behavioral demands explicit.

The argument–adjunct manipulation supplies a defined linguistic context for these measurements. Arguments are selected by a predicate and prototypically express core event participants, whereas adjuncts enrich an event with less tightly selected information [12,13]. Faster affected-subject judgments after argument completions are consistent with closer alignment between the sentence’s core event structure and the question being asked. Direct-object nouns and manner adverbs also differ in grammatical category and uncontrolled lexical and acoustic properties. The result might therefore concern this manipulation and response demand, rather than a general argument-processing advantage.

The behavioral association raises a related question, does preserved intrinsic organization become more consequential when semantic integration is less externally constrained? The consistent cross-fitting directions motivate this possibility, but the corrected estimate and the moderation test do not yet establish it. A larger study sampling independently normed degrees of lexical selection, core-event status, and semantic fit could test that account across a broader argumenthood continuum [12].

The study’s scope is set by its 14-person main cohort and within-session sensor-space design. Repeated trials improve each participant’s estimates but do not replace independent participant replication. Cross-fitting separates task observations without providing out-of-sample prediction. The ERP contrast has limited individual reliability and overlaps the response period, while the behavioral inference depends partly on the reference pool. CSD reduces reference and broad-field sensitivity, but head anatomy and cap geometry can still contribute to repeatable topographies. These limitations favor replication with repeated sessions, a larger cohort, and linguistic materials that separate completion type from lexical and acoustic properties.

Person-specific intrinsic-timescale topographies persist from rest into sentence processing, and absolute task timescale relates to the late-positive response to semantic anomaly. The presented results locate these findings in the same individuals. Behavioral estimates provide a focused target for replication, linking the preservation of temporal organization to the cost of semantic decisions.

## References

1. Northoff G, Qin P, Nakao T. Rest-stimulus interaction in the brain: a review. Trends Neurosci. 2010;33:277–284. doi:10.1016/j.tins.2010.02.006.

2. Golesorkhi M, Gomez-Pilar J, Zilio F, Berberian N, Wolff A, Yagoub MCE, et al. The brain and its time: intrinsic neural timescales are key for input processing. Commun Biol. 2021;4:970. doi:10.1038/s42003-021-02483-6.

3. Wolff A, Berberian N, Golesorkhi M, Gomez-Pilar J, Zilio F, Northoff G. Intrinsic neural timescales: temporal integration and segregation. Trends Cogn Sci. 2022;26:159–173. doi:10.1016/j.tics.2021.11.007.

4. Northoff G, Wainio-Theberge S, Evers K. Spatiotemporal neuroscience - what it is and why we need it. Phys Life Rev. 2020;33:78–87. doi:10.1016/j.plrev.2020.06.005.

5. Northoff G, Wainio-Theberge S, Evers K. Is temporo-spatial dynamics the “common currency” of brain and mind? Phys Life Rev. 2020;33:34–54. doi:10.1016/j.plrev.2019.05.002.

6. Murray JD, Bernacchia A, Freedman DJ, Romo R, Wallis JD, Cai X, et al. A hierarchy of intrinsic timescales across primate cortex. Nat Neurosci. 2014;17:1661–1663. doi:10.1038/nn.3862.

7. Honey CJ, Thesen T, Donner TH, Silbert LJ, Carlson CE, Devinsky O, et al. Slow cortical dynamics and the accumulation of information over long timescales. Neuron. 2012;76:423–434. doi:10.1016/j.neuron.2012.08.011.

8. Çatal Y, Keskin K, Wolman A, Klar P, Smith D, Northoff G. Flexibility of intrinsic neural timescales during distinct behavioral states. Commun Biol. 2024;7:1667. doi:10.1038/s42003-024-07349-1.

9. Tavor I, Parker Jones O, Mars RB, Smith SM, Behrens TE, Jbabdi S. Task-free MRI predicts individual differences in brain activity during task performance. Science. 2016;352:216–220. doi:10.1126/science.aad8127.

10. Finn ES, Shen X, Scheinost D, Rosenberg MD, Huang J, Chun MM, et al. Functional connectome fingerprinting: identifying individuals using patterns of brain connectivity. Nat Neurosci. 2015;18:1664–1671. doi:10.1038/nn.4135.

11. Gratton C, Laumann TO, Nielsen AN, Greene DJ, Gordon EM, Gilmore AW, et al. Functional brain networks are dominated by stable group and individual factors, not cognitive or daily variation. Neuron. 2018;98:439–452.e5. doi:10.1016/j.neuron.2018.03.035.

12. Toivonen I. Arguments and adjuncts across levels. In: Butt M, Findlay JY, Toivonen I, editors. Proceedings of the LFG’21 Conference. Stanford: CSLI Publications; 2021. p. 306–331.

13. Van Luven K, Toivonen I. The obligatoriness of arguments. Lang Linguist Compass. 2024;18:e12511. doi:10.1111/lnc3.12511.

14. Tutunjian D, Boland JE. Do we need a distinction between arguments and adjuncts? Evidence from psycholinguistic studies of comprehension. Lang Linguist Compass. 2008;2:631–646. doi:10.1111/j.1749-818X.2008.00071.x.

15. Koenig JP, Mauner G, Bienvenue B. Arguments for adjuncts. Cognition. 2003;89:67–103. doi:10.1016/S0010-0277(03)00082-9.

16. Kennison SM. Comprehending noun phrase arguments and adjuncts. J Psycholinguist Res. 2002;31:65–81. doi:10.1023/A:1014328321363.

17. Speer SR, Clifton C Jr. Plausibility and argument structure in sentence comprehension. Mem Cognit. 1998;26:965–978. doi:10.3758/BF03201177.

18. Smith D, Wolff A, Wolman A, Ignaszewski J, Northoff G. Temporal continuity of self: long autocorrelation windows mediate self-specificity. Neuroimage. 2022;257:119305. doi:10.1016/j.neuroimage.2022.119305.

19. Perrin F, Pernier J, Bertrand O, Echallier JF. Spherical splines for scalp potential and current density mapping. Electroencephalogr Clin Neurophysiol. 1989;72:184–187. doi:10.1016/0013-4694(89)90180-6.

20. Kayser J, Tenke CE. Principal components analysis of Laplacian waveforms as a generic method for identifying ERP generator patterns: I. Evaluation with auditory oddball tasks. Clin Neurophysiol. 2006;117:348–368. doi:10.1016/j.clinph.2005.08.034.

21. Nichols TE, Holmes AP. Nonparametric permutation tests for functional neuroimaging: a primer with examples. Hum Brain Mapp. 2002;15:1–25. doi:10.1002/hbm.1058.

22. Westfall PH, Young SS. Resampling-based multiple testing: examples and methods for p-value adjustment. New York: Wiley; 1993.

23. Benjamini Y, Hochberg Y. Controlling the false discovery rate: a practical and powerful approach to multiple testing. J R Stat Soc B. 1995;57:289–300. doi:10.1111/j.2517-6161.1995.tb02031.x.

24. Northoff G, Wolman A, Zhang J. Brain dynamics shape cognition—Spatiotemporal Neuroscience. Phys Life Rev. 2025;54:173–201. doi:10.1016/j.plrev.2025.07.009.

25. Wainio-Theberge S, Wolff A, Northoff G. Dynamic relationships between spontaneous and evoked electrophysiological activity. Commun Biol. 2021;4:741. doi:10.1038/s42003-021-02240-9.

26. Kutas M, Federmeier KD. Thirty years and counting: finding meaning in the N400 component of the event-related brain potential. Annu Rev Psychol. 2011;62:621–647. doi:10.1146/annurev.psych.093008.131123.

27. Kuperberg GR. Neural mechanisms of language comprehension: challenges to syntax. Brain Res. 2007;1146:23–49. doi:10.1016/j.brainres.2006.12.063.

28. Brouwer H, Fitz H, Hoeks J. Getting real about semantic illusions: rethinking the functional role of the P600 in language comprehension. Brain Res. 2012;1446:127–143. doi:10.1016/j.brainres.2012.01.055.

29. Friederici AD. The brain basis of language processing: from structure to function. Physiol Rev. 2011;91:1357–1392. doi:10.1152/physrev.00006.2011.

30. Çatal Y, Keskin K, Wolman A, Buccellato A, Northoff G. Intrinsic neural timescales relate to event-related activity—key role for intracolumnar connections. J Neurosci. 2025;45:e0896252025. doi:10.1523/JNEUROSCI.0896-25.2025.

31. Chuipka N, Smy T, Northoff G. From neural activity to behavioral engagement: temporal dynamics as their “common currency” during music. Neuroimage. 2025;312:121209. doi:10.1016/j.neuroimage.2025.121209.

