## Supplementary Materials for "Person-specific intrinsic-timescale topographies persist from rest into sentence processing"

The main manuscript centers on the same 14 participants with eyes-open rest EEG, task timescales, eligible ERP effects, and behavior. This supplement preserves the wider-cohort checks, complete multiplicity families, measurement diagnostics, and additional analyses. Supporting samples are identified separately and are not independent replication cohorts.

#### S1 Cohort definition and supporting samples

The main cohort comprised of participants that met the fixed eyes-open rest–task criteria and the all-block and split-half ERP thresholds. Of the 15 eyes-open rest–task participants, one lacked the required even-half ERP coverage and was therefore excluded from the main cohort. The remaining exclusions reflect measurement eligibility rather than splitting an otherwise complete sample.

**Table S1 | Roles of the available cohorts**

| Cohort | N | Role |
|---|---|---|
| Main common cohort | 14 | All main figures, ERP predictors and behavioral tests; 14-person mismatch pool |
| Task behavior | 21 | Supporting group behavioral effects |
| Pooled ERP effect | 18 | Supporting late-positive group response |
| Eyes-open rest–task | 15 | Original fingerprint, channel controls and behavioral reference pool |
| Eyes-closed rest–task | 13 | Supporting separate-state fingerprint and behavioral estimate |
| Matched resting states | 11 | Main-cohort subset with both resting states |
| Anomalous structure contrast | 19 | Supporting adjunct-anomalous minus argument-anomalous ERP comparison |

All samples arise from the 22-person data archive. Different endpoint requirements account for the additional eligible observations. The 19-person ERP comparison requires only anomalous trials, whereas the pooled effect also requires plausible trials.

#### S2 Spatial and reference controls

The main 14 × 14 fingerprint matrices recompute each person's mismatch mean from the other 13 people. The larger-sample checks below retain their original eligible cohorts. Group-centering subtracts the cohort-average channel pattern separately from rest and task. Channel exclusion omits channels flagged before interpolation in either recording; the task mask is conservative across blocks.

In the 15-person eyes-open sample, the average-reference own-person advantage was $\Delta z = 0.469$, with 66.7% identification. Group-centering increased the advantage to 0.569 and identification to 86.7%. Excluding pre-interpolation bad channels retained $\Delta z = 0.467$ and 53.3% identification. The CSD own-person advantage was positive in every task block. These tests support preservation of an individual configuration, while the same-session sensor-space design retains possible effects of head anatomy and electrode geometry.

In the full 13-person eyes-closed sample, rest provided a separate-state validation. Own-person similarity again exceeded mismatch (own z = 0.536, mismatch = 0.195, advantage = 0.341; dz = 1.571; $t(12) = 5.67$, $P = 1.05 \times 10^{-4}$; permutation $P = 5.00 \times 10^{-5}$). Eight of 13 participants were identified correctly (61.5% versus 7.7% chance, $P = 5.00 \times 10^{-5}$). Group-centering yielded a 0.414 advantage and 76.9% identification; excluding pre-interpolation bad channels yielded a 0.360 advantage and 53.8% identification. The recurrence supports a within-session person-specific configuration across resting states.

**Table S2 | Fingerprints in the same 11 participants with both resting states**

| Resting state | Reference | Advantage Δz | Identity | Continuity P |
|---|---|---|---|---|
| eyes closed | Average | 0.340 | 7/11 | 5e-06 |
| eyes closed | Cz | 0.333 | 7/11 | 5e-06 |
| eyes closed | CSD | 0.348 | 7/11 | 5e-06 |
| eyes open | Average | 0.476 | 9/11 | 5e-06 |
| eyes open | Cz | 0.362 | 6/11 | 5e-06 |
| eyes open | CSD | 0.571 | 9/11 | 5e-06 |

Each restricted matrix uses the other 10 participants for mismatch comparisons and 1/11 chance identification. Results use 200,000 participant-label permutations. Figure S1 shows the full 15-person eyes-open and 13-person eyes-closed CSD analyses.

### S3 Complete behavioral inference family

All six behavioral endpoints are retained in the familywise correction, including those not emphasized in the main narrative. Every row in Table S3 uses the main 14 participants and a mismatch pool restricted to those same people. A single permutation reassigns both behavioral halves together across participants. Pointwise bootstrap intervals are not familywise confidence intervals.

**Table S3 | Six behavioral associations in the main cohort**

| Endpoint | ρ | Nominal P | MinP P | Holm P |
|---|---|---|---|---|
| Affected anomaly | 0.547 | 0.009 | 0.054 | 0.056 |
| Affected structure | -0.073 | 0.766 | 1.000 | 1.000 |
| Affected interaction | -0.422 | 0.075 | 0.369 | 0.375 |
| Understood anomaly | 0.024 | 0.928 | 1.000 | 1.000 |
| Understood structure | 0.204 | 0.423 | 0.958 | 1.000 |
| Understood interaction | -0.040 | 0.872 | 1.000 | 1.000 |

ρ is the mean of the two opposite-half Spearman correlations. MinP denotes Westfall–Young single-step familywise correction. Structure is argument minus adjunct; interaction is structure × anomaly. The anomaly association has nominal P = .0093 but does not meet the familywise .05 threshold.

### S4 Behavioral cohort and estimator sensitivities

Adopting the common cohort changes two distinct things: which participants enter the association, and which resting topographies define the mismatch baseline. The earlier 14-person sensitivity changed only the inferential sample and retained the original 15-person resting pool. The main analysis now restricts both to 14 people. The corresponding changes in the estimate and corrected P value are shown together below. No threshold, endpoint, or multiplicity-family member was changed to obtain either result.

**Table S4 | Effect of cohort and reference-pool definition**

| Analysis | Inferential N | Rest pool N | ρ | Familywise P |
|---|---|---|---|---|
| Main cross-fitted analysis | 14 | 14 | 0.547 | 0.054 |
| Earlier common-cohort sensitivity | 14 | 15 | 0.574 | 0.031 |
| Original eyes-open analysis | 15 | 15 | 0.550 | 0.036 |
| All-block main-cohort sensitivity | 14 | 14 | 0.688 | 0.045 |

All-block estimates reuse task observations for the neural and behavioral measures and therefore do not replace opposite-half inference. The original 15-person result is retained from the v1.1.8 behavioral-stage export. These are dependent sensitivity analyses, not independent replications.

**Table S5 | Alternative behavioral estimators in the same 14-person cohort**

| Estimator | ρ | Pointwise 95% CI | Nominal P |
|---|---|---|---|
| Adjusted log RT | 0.547 | 0.197 to 0.764 | 0.009 |
| Cellwise log median | 0.446 | 0.108 to 0.665 | 0.033 |
| 10% trimmed log mean | 0.527 | 0.160 to 0.763 | 0.013 |
| Raw RT median | 0.451 | 0.089 to 0.694 | 0.037 |

These estimates use the same 14-person resting pool. Nominal P values are sensitivity summaries and are not separately presented as confirmatory findings. The primary six-endpoint correction remains Table S3.

Within the main cohort, the average anomaly coefficient was 0.087 (95% CI −0.004 to 0.178; P = 0.059), corresponding to 9.1% slowing. The structure-by-anomaly group coefficient was −0.085 (95% CI −0.189 to 0.019; P = 0.102). Simple anomaly costs were descriptively 13.8% for adjuncts and 4.6% for arguments. These decompositions do not establish a between-structure difference.

In the full 21-person behavioral sample, arguments were judged 15.5% faster (95% CI 7.6–22.7%; P = 0.000825), and the average anomaly cost was 8.1% (P = 0.028). In the original 15-person continuity analysis, the structure-specific association estimates were ρ = 0.598 for adjuncts and 0.141 for arguments; formal moderation did not survive correction (familywise P = 0.243). Figure S4 retains those wider-cohort displays as support, with their sample sizes explicit.

The eyes-closed behavioral association was small in its full eligible sample (N = 13; ρ = 0.077; P = 0.775). Among the 11 participants with both resting states, estimates were ρ = 0.545 for eyes-open rest (95% CI 0.174–0.804) and ρ = 0.177 for eyes-closed rest (95% CI −0.320 to 0.659). These earlier matched-state behavioral checks retained the original state-specific reference pools. The estimates do not establish state specificity.

#### S5 ERP diagnostics and interpretation

The main cohort shows a group late positivity but limited reliability of the individual anomaly contrast. Table S6 reports both Pearson and rank reproducibility so the group effect is not mistaken for a stable individual phenotype.

**Table S6 | Split-half measurement reproducibility in the main cohort**

| Measure | N | Pearson r | Spearman ρ |
|---|---|---|---|
| CSD continuity | 14 | 0.965 | 0.930 |
| Anomaly-cost coefficient | 14 | 0.836 | 0.336 |
| Absolute task timescale | 14 | 0.988 | 0.925 |
| Late-positive ERP contrast | 14 | 0.206 | 0.156 |

Both halves contain the same participants. Continuity uses the 14-person resting reference pool. These are reliability estimates, not evidence of independent replication.

Across the 18 ERP-eligible participants, the pooled anomaly effect was +0.530 µV (95% CI 0.284–0.775; dz = 1.072; $P = 0.000286$). The 17 participants eligible in both task halves had Pearson split-half reliability $r = 0.025$. In this 18-person group, 62.4% of retained trial response markers preceded 900 ms, compared with 59.6% in the main 14-person cohort (median latency 0.806 s). Figure S2 preserves the broader-cohort waveform, reliability, and timing diagnostics.

The response-safe sensitivity retained trials whose marker followed the 900-ms window end by at least 200 ms. It retained 11 participants and gave +0.255 µV (95% CI −0.702 to 1.211; $P = 0.566$). The result is imprecise and does not isolate a response-free P600 mechanism. The anomalous-adjunct minus anomalous-argument contrast required at least 40 accepted trials in each anomalous cell, without requiring plausible trials. Across its 19 eligible participants, it was +0.138 µV (95% CI −0.411 to 0.687; $P = 0.603$). It does not establish a structure-specific ERP effect.

Çatal and colleagues [30] reported a positive relation between a resting half-decay timescale and MEG event-related field magnitude. The present test uses an EEG zero-crossing timescale and a signed anomalous-minus-plausible voltage contrast. Its negative task-timescale association therefore addresses a different measurement and does not directly replicate the prior effect or establish an intracolumnar mechanism. The resting and task predictors are not mutually adjusted, and their significance levels do not test a difference between associations.

#### S6 Temporal variability in timescales and response times

This additional three-test family uses the same 14 participants as the main cohort. It concerns dispersion across repeated windows and trials, rather than the absolute-timescale predictors or preservation of spatial topography.

To test whether variability in intrinsic timescales related to behavioral variability [8], we used the same accepted 4-s CSD windows as the topographic analysis. For each channel, temporal variability was the sample SD of finite zero-crossing estimates across windows; participant variability was the median of eligible channel SDs. Rest windows did not overlap. Task windows remained locked to critical-word onset and could overlap in time. Eligibility required ≥80% finite estimates per channel, ≥30 eligible channels, ≥30 resting windows (≥15 per reliability half), and ≥40 task windows per analyzed split, with all four target conditions represented. The existing minimum of four accepted trials per block × verb × condition was retained. A sensitivity removed each block × verb × structure × plausibility cell mean before calculating a pooled within-cell SD.

Behavioral variability was the residual SD of log affected-subject response times after adjustment for anomaly, structure, their interaction, verb, and block order, calculated as the square root of residual sum of squares divided by residual degrees of freedom. Each model required full rank and ≥20 trials. Three associations formed a separate Holm-corrected family on the same 14 eligible participants: resting timescale SD with response-time SD; opposite-half task timescale SD with response-time SD; and resting with task timescale SD. Inference used 200,000 participant-label permutations and 5,000 paired participant bootstrap resamples. Reliability used interleaved rest segments or the task-block halves. These SDs describe dispersion over repeated windows and trials, not moment-by-moment neural–response coupling.

Temporal dispersion of the CSD timescale estimates was reproducible across halves: Spearman $\rho = 0.907$ for resting SD ($N = 15$) and $\rho = 0.969$ for task SD ($N = 14$). Residual log-response-time SD was also reproducible ($\rho = 0.929$, $N = 15$). Nevertheless, within the common 14-person test family, resting timescale SD was not reliably associated with response-time SD ($\rho = 0.077$, 95% CI −0.582 to 0.685, Holm-adjusted $P = 1.00$), nor was opposite-half task timescale SD ($\rho = -0.002$, 95% CI −0.616 to 0.578, Holm-adjusted $P = 1.00$; Fig. S3). Removing task-cell mean differences gave a similar result ($\rho = 0.015$, nominal $P = 0.960$).

Resting and task timescale SDs showed a positive point estimate ($\rho = 0.604$, 95% CI 0.022–0.911, nominal $P = 0.025$), but the association did not survive correction across the three tests (Holm-adjusted $P = 0.075$). The intervals are pointwise rather than multiplicity-adjusted. Thus, the variability analysis did not establish an additional neural–behavioral association, and its broad intervals leave moderate associations unresolved. The complete three-test family is summarized in Table S7.

**Table S7 | Temporal-variability family in the main cohort**

| Association | ρ | Pointwise 95% CI | Holm P |
|---|---|---|---|
| Rest variability–RT variability | 0.077 | -0.582 to 0.685 | 1.000 |
| Task variability–RT variability | -0.002 | -0.616 to 0.578 | 1.000 |
| Rest–task variability | 0.604 | 0.022 to 0.911 | 0.075 |

All rows have N = 14. Task–RT inference pairs opposite task halves. Holm correction includes all three tests. The pointwise interval for the rest–task estimate is not a multiplicity-adjusted interval.

Chuipka and colleagues linked EEG and continuous engagement-rating dynamics during music using dynamic median frequency [31]. The present dispersion analysis measures a different form of neural–behavioral coupling and provides no additional corrected association. Window-estimation noise, unequal accepted-window counts, and overlapping task windows may contribute to the neural SDs. The estimates should not be equated with direct neural flexibility or moment-by-moment neural–response coupling.

#### Supplementary references

Reference numbers correspond to the main manuscript.

8. Çatal Y, Keskin K, Wolman A, Klar P, Smith D, Northoff G. Flexibility of intrinsic neural timescales during distinct behavioral states. Commun Biol. 2024;7:1667. doi:10.1038/s42003-024-07349-1.

30. Çatal Y, Keskin K, Wolman A, Buccellato A, Northoff G. Intrinsic neural timescales relate to event-related activity—key role for intracolumnar connections. J Neurosci. 2025;45:e0896252025. doi:10.1523/JNEUROSCI.0896-25.2025.

31. Chuipka N, Smy T, Northoff G. From neural activity to behavioral engagement: temporal dynamics as their “common currency” during music. Neuroimage. 2025;312:121209. doi:10.1016/j.neuroimage.2025.121209.

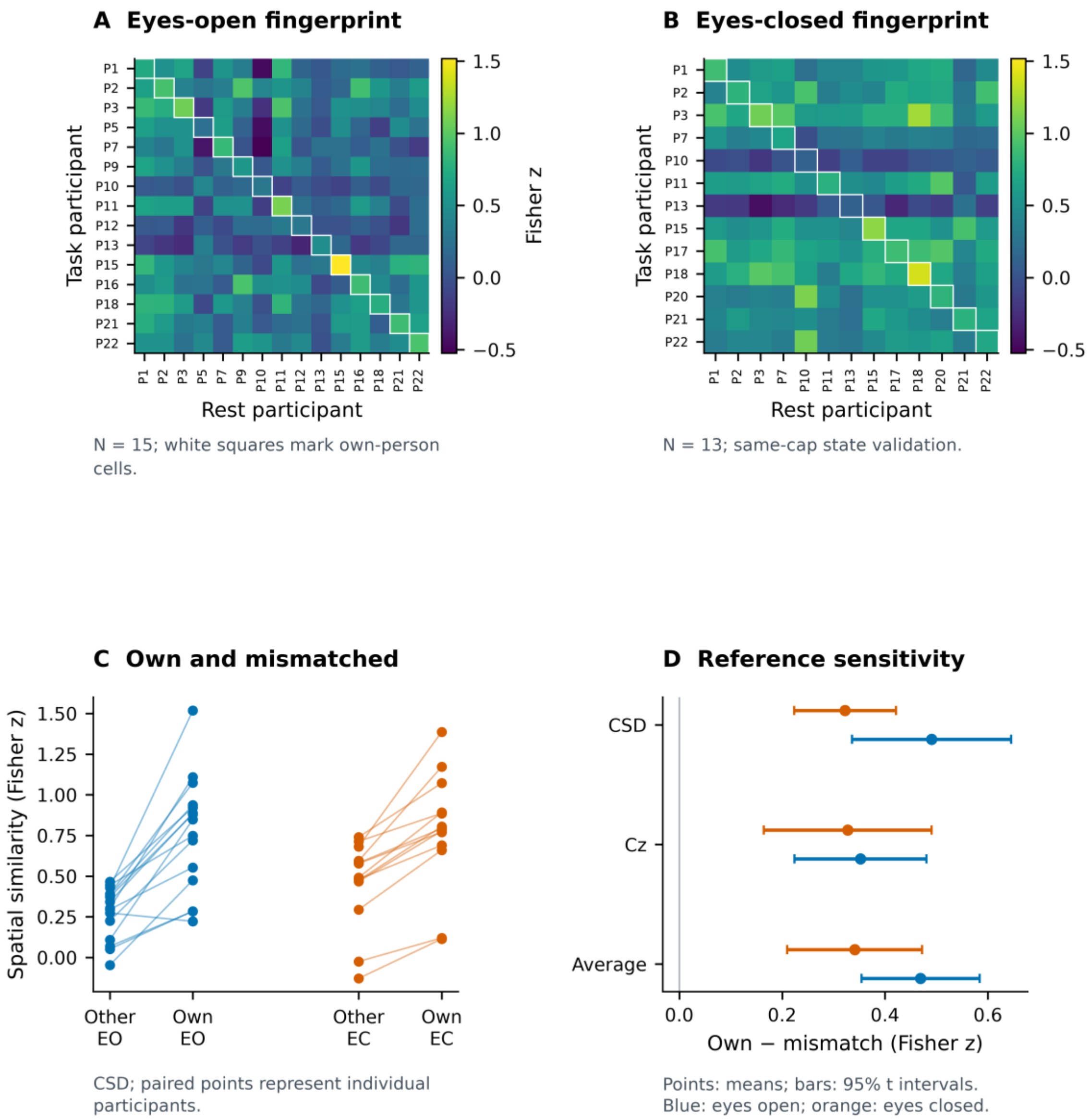


Figure S1. Fingerprints in the larger eligible resting-state cohorts. (A,B) CSD task–rest matrices for eyes-open rest (N = 15) and eyes-closed rest (N = 13). (C) Paired own-person and mismatched similarities. (D) Reference-sensitivity means with 95% t intervals. These supporting cohorts differ from the fixed 14-person main cohort and are not independent samples. All displayed continuity permutation P values are $5.00 \times 10^{-5}$.

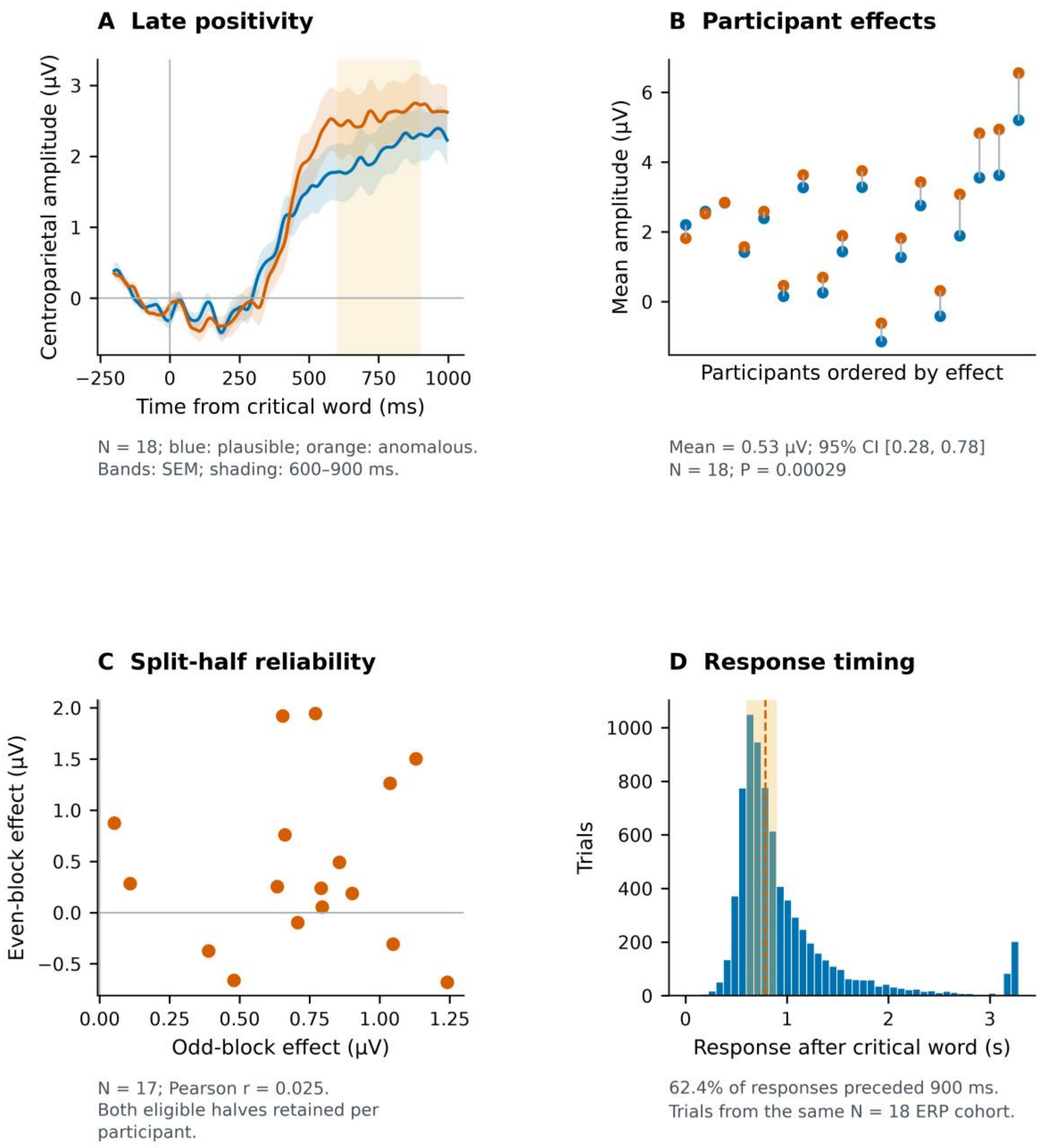


Figure S2. ERP diagnostics in the larger ERP-eligible cohort. (A) Mean ± SEM waveforms and (B) paired condition amplitudes for N = 18. (C) Split-half effects in the 17 participants meeting both-half criteria (Pearson r = 0.025). (D) Response timing in the 18-person ERP cohort; 62.4% preceded 900 ms. These supporting counts differ from the N = 14 main-cohort waveform and timing results.

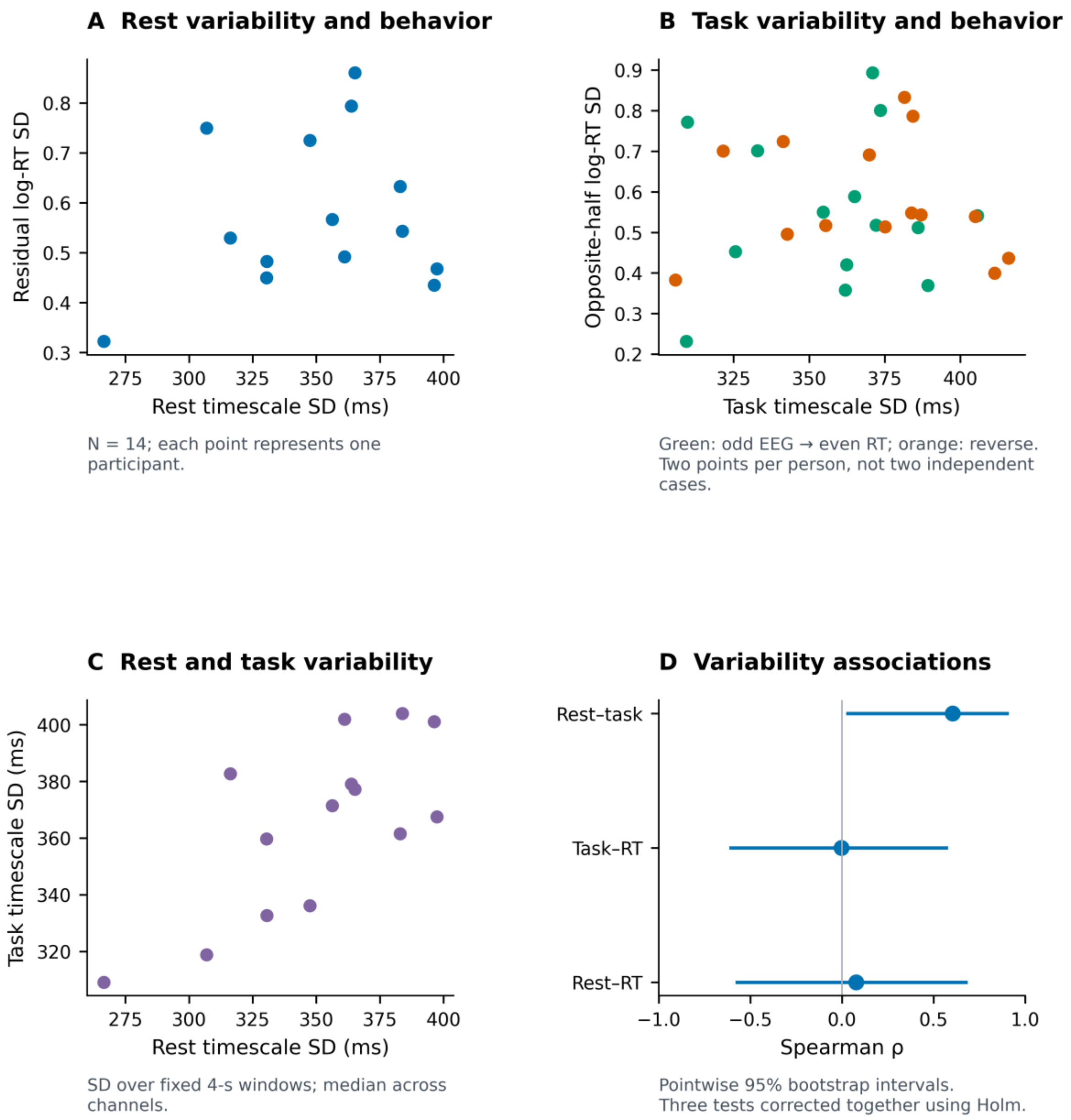


Figure S3. Temporal variability in the same 14 participants as the main cohort. (A) Resting timescale SD and residual log-RT SD. (B) Opposite-half task timescale SD and RT SD; two points per person do not increase the inferential N. (C) Rest and task SDs. (D) Associations with pointwise 95% bootstrap intervals. Holm-adjusted P values are 1.00, 1.00, and 0.075. Neural SD is calculated across accepted windows within channels, then summarized by the channel median.

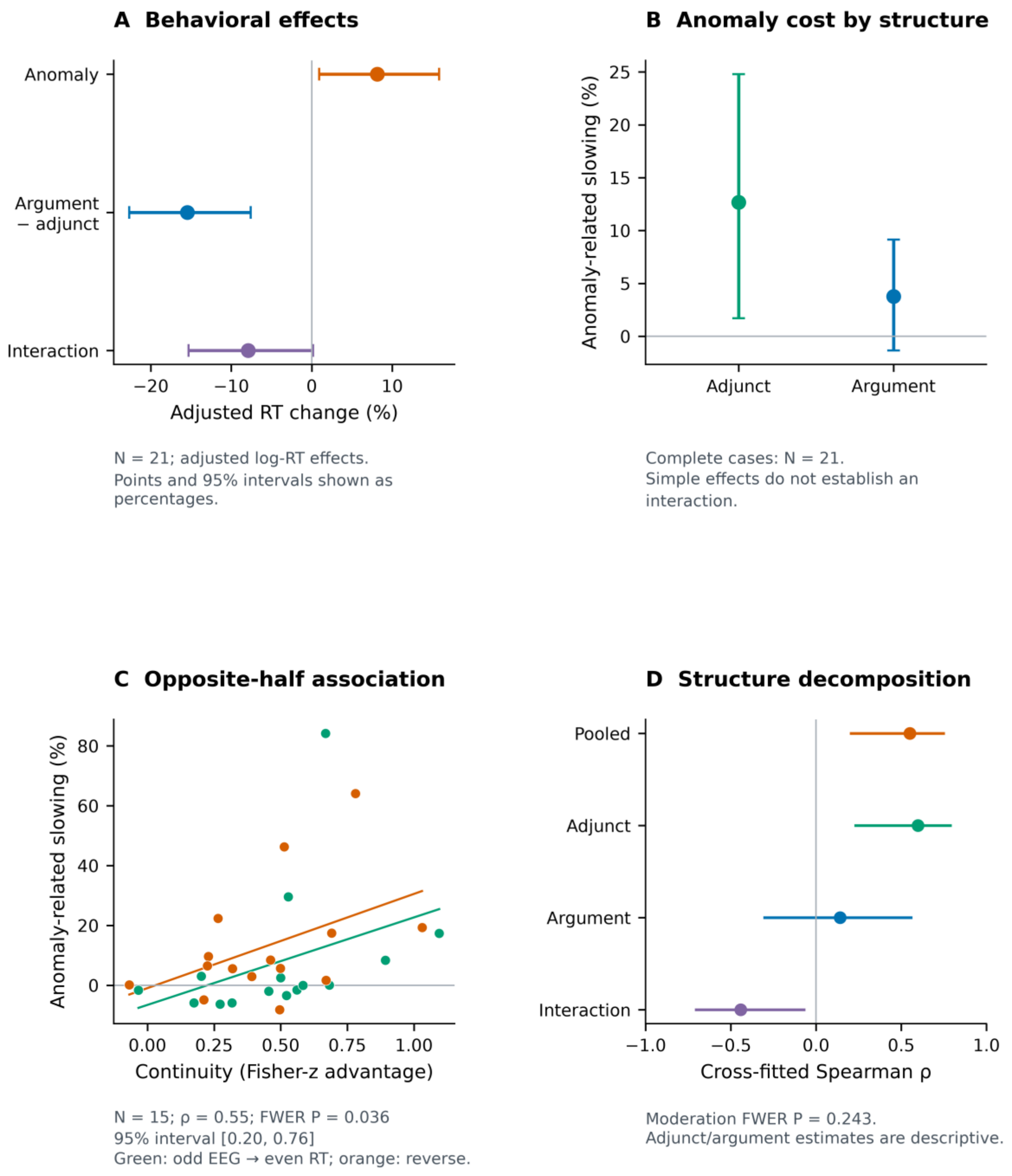


Figure S4. Behavioral findings in the larger supporting cohorts. (A,B) Group and structure-specific behavioral contrasts use N = 21. (C,D) Continuity–behavior and structure-decomposition analyses use N = 15 and the original 15-person resting pool. The original pooled familywise P is 0.036; moderation P is 0.243. These displays document the earlier broader-cohort results. Main-cohort inference with N = 14 and a 14-person resting pool is shown in main Figure 4 and Supplementary Table S3.
